# RepCOOL: Computational Drug Repositioning Via Integrating Heterogeneous Biological Networks

**DOI:** 10.1101/817882

**Authors:** Ghazale Fahimian, Javad Zahiri, Seyed Sh. Arab, Reza H. Sajedi

## Abstract

**Background:** It often takes more than 10 years and costs more than one billion dollars to develop a new drug for a disease and bring it to the market. Drug repositioning can significantly reduce costs and times in drug development. Recently, computational drug repositioning attracted a considerable amount of attention among researchers, and a plethora of computational drug repositioning methods have been proposed.

**Methods:** In this study, we propose a novel network-based method, named RepCOOL, for drug repositioning. RepCOOL integrates various heterogeneous biological networks to suggest new drug candidates for a given disease.

**Results:** The proposed method showed a promising performance on benchmark datasets via rigorous cross-validation. Final drug repositioning model has been built based on random forest classifier, after examining various machine learning algorithms. Finally, in a case study, four FDA approved drugs were suggested for breast cancer stage II.

**Conclusion:** Results show the strength of the proposed method in detecting true drug-disease relationships. RepCOOL suggested four new drugs for breast cancer stage II namely Doxorubicin, Paclitaxel, Trastuzumab and Tamoxifen.

## Background

Drug research and development is a time-consuming, expensive and complicated process. Previous research reported that it often takes 10–15 years and 0.8–1.5 billion dollars to develop a new drug and bring it to the market [1]. Although such a huge amount of time and money is expanding in this industry, the number of new FDA-approved drugs annually remains low. So, in light of these challenges, finding a new use for an existing drug, which is known as drug repositioning or drug repurposing, has been proposed as a solution for such problem. The goal of drug repositioning is to identify new indications for existing drugs. The result of using such approaches can reduce the overall cost of commercialization, and also eliminate the delay between drug discovery and availability. In comparison to the traditional drug repositioning which relies on clinical discoveries, computational drug repositioning methods can simplify the drug development timeline[2–6].

In recent years, different approaches were exploited for repurposing drugs, including network, text mining, machine learning and semantic inference based approaches. Recently, network-based approach attracted more attention and was widely used in computational drug repositioning due to the capability of using ever-increasing large scale biological datasets such as genetic, pharmacogenomics, clinical and chemical data [2, 5, 7–14].

In this study, we have proposed a network-based approach for drug repositioning. Our method, namely RepCOOL, integrated various heterogeneous biological networks to obtain new drug-disease associations. The proposed method showed a satisfactory performance in detecting drug-disease associations via stringent assessment procedures. Eventually, four new drugs were suggested for breast cancer.

## 2. Method

Figure 1 depicts a schematic flowchart of the proposed drug repositioning method. Detailed descriptions for each step were provided in the following subsections.

**Figure 1.**
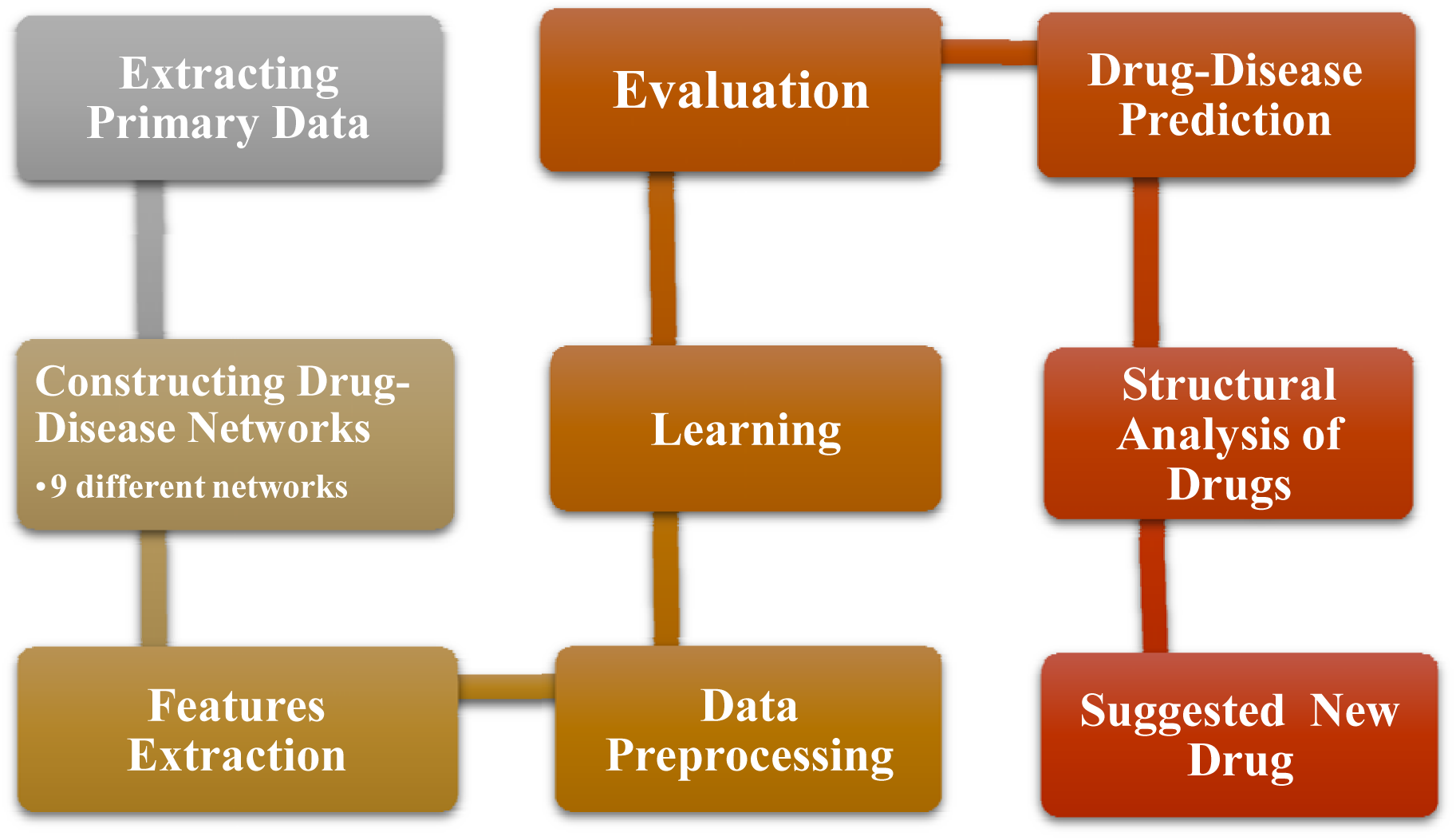
Schematic flowchart of the proposed drug repositioning method.

### 2.1 Data sources

We have constructed nine different drug-disease association networks using six primary networks that were constructed based on the publicly available database (Table 1). These six network can be categorized into four different groups according to their types of nodes: drug-gene interaction network (DRGN), disease-gene interaction network (DIGN), protein-protein interaction network (PPIN) and gene co-expression network (GCN).

**Table 1.**
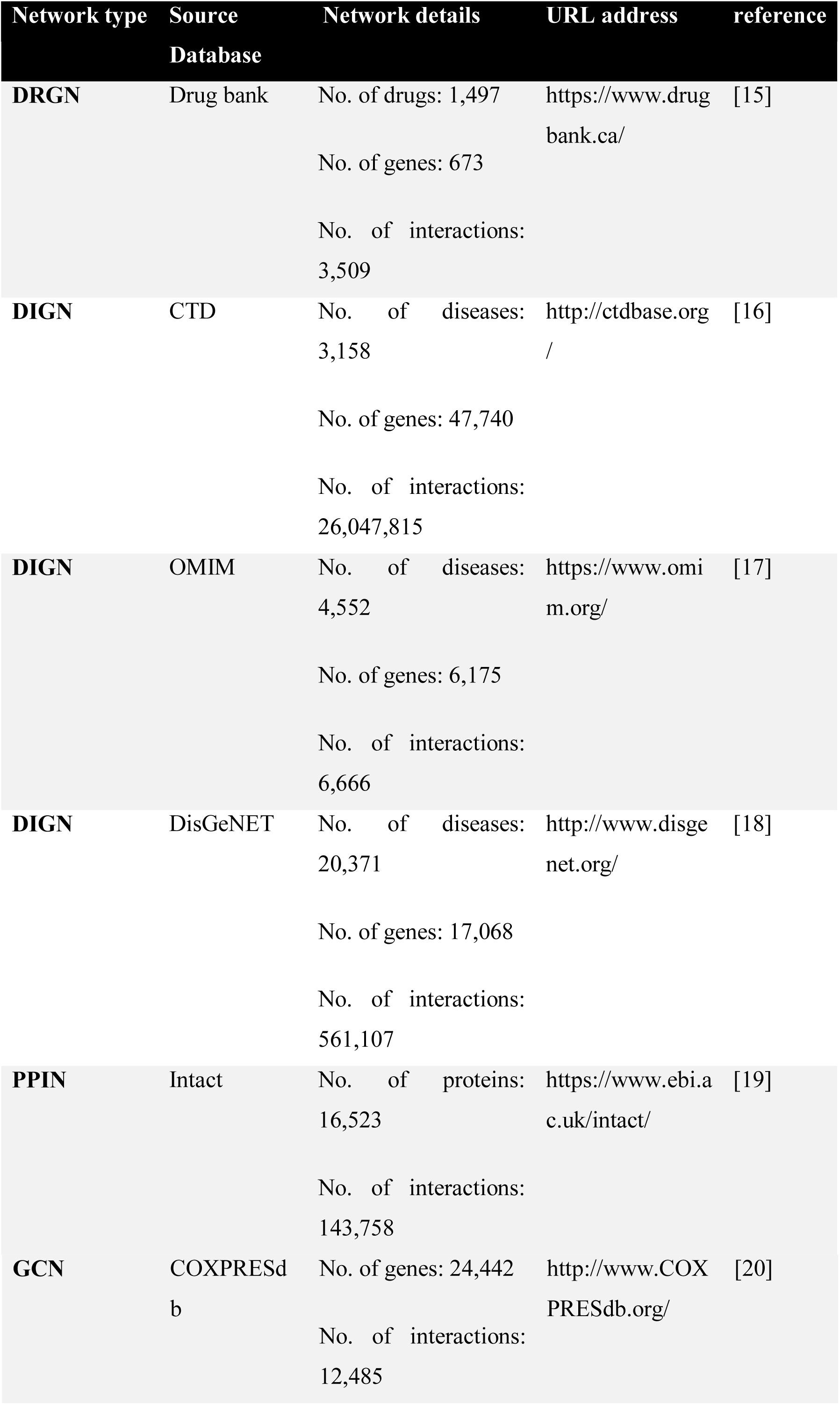
Primary data sources for drug-disease network reconstruction.

#### Drug-gene interaction network

We used DrugBank [15] database to construct DRGN network. DrugBank provides comprehensive information about approved and investigational drugs, including UMLS-mapped approved indications. This network includes 3,509 interactions between 1,497 drugs and 673 genes.

#### Disease-gene interaction network

We used three databases for constructing three different disease-gene interaction networks (Table 1): The Comparative Toxic genomics Database (CTD) [16], Online Mendelian Inheritance in Man (OMIM) [17] and DisGeNET[18]. CTD contains manually curated information about gene-disease relationships with focus on understanding the effects of environmental chemicals on human health and included more than 26 million gene–disease associations (GDAs), between 47,740 genes and 3,158 diseases. OMIM (Online Mendelian Inheritance in Man) is a complete collection of human genes and genetic phenotypes that is updated daily. OMIM includes 6,666 gene-phenotype associations between 6,175 phenotypes and 4,552 genes. The DisGeNET database integrates human gene-disease associations from various expert curated databases and text-mining derived associations including Mendelian, complex and environmental diseases[18]. This network included 561,107 GDAs, between 17,068 genes and 20,371 diseases, disorders, traits, and clinical or abnormal human phenotypes.

#### Protein-protein interaction network

We extracted protein-protein interaction (PPI) information from IntAct database[19]. IntAct provides a freely available database system and analysis tools for molecular interaction data. This network has 16,523 proteins and 143,738 protein-protein interactions.

#### Gene co-expression network

We have constructed gene co-expression network (GCN) using COXPRESdb database[20]. This database measured the similarity of gene expression patterns during several conditions such as disease states tissue types. COXPRESdb includes co-expression relationships for multiple animal species and is freely available on http://coxpresdb.jp/. The obtained GCN includes 12,485 interactions and 24,442 genes.

### 2.2 Reconstructing new drug-disease networks via merging heterogeneous networks

We have reconstructed nine new drug-disease networks using six primary networks. Figure 2 shows a schematic of the reconstruction of these networks. These nine networks have more than 9,400,000 drug-disease associations in total. Table 2 shows more details about these new drug-disease networks. One drug-disease interaction may be generated more than once in each network merging. So, the number of occurrences of a drug-disease interaction is considered as the weight of that interaction.

**Table 2.** Reconstructed drug-disease networks

| **Networks** | **Number of Drug** | **Number of Disease** | **Drug-Disease association** |
| --- | --- | --- | --- |
| **Net1** | 1,337 | 5,854 | 4,129,617 |
| **Net2** | 1,333 | 8,540 | 397,108 |
| **Net3** | 1,191 | 10,858 | 741,819 |
| **Net4** | 1,208 | 11,934 | 8,256,300 |
| **Net5** | 164 | 2,240 | 82,407 |
| **Net6** | 239 | 2,306 | 92,299 |
| **Net7** | 94 | 2,200 | 151,267 |
| **Net8** | 21 | 1,013 | 329 |
| **Net9** | 17 | 468 | 812 |

**Figure 2.**
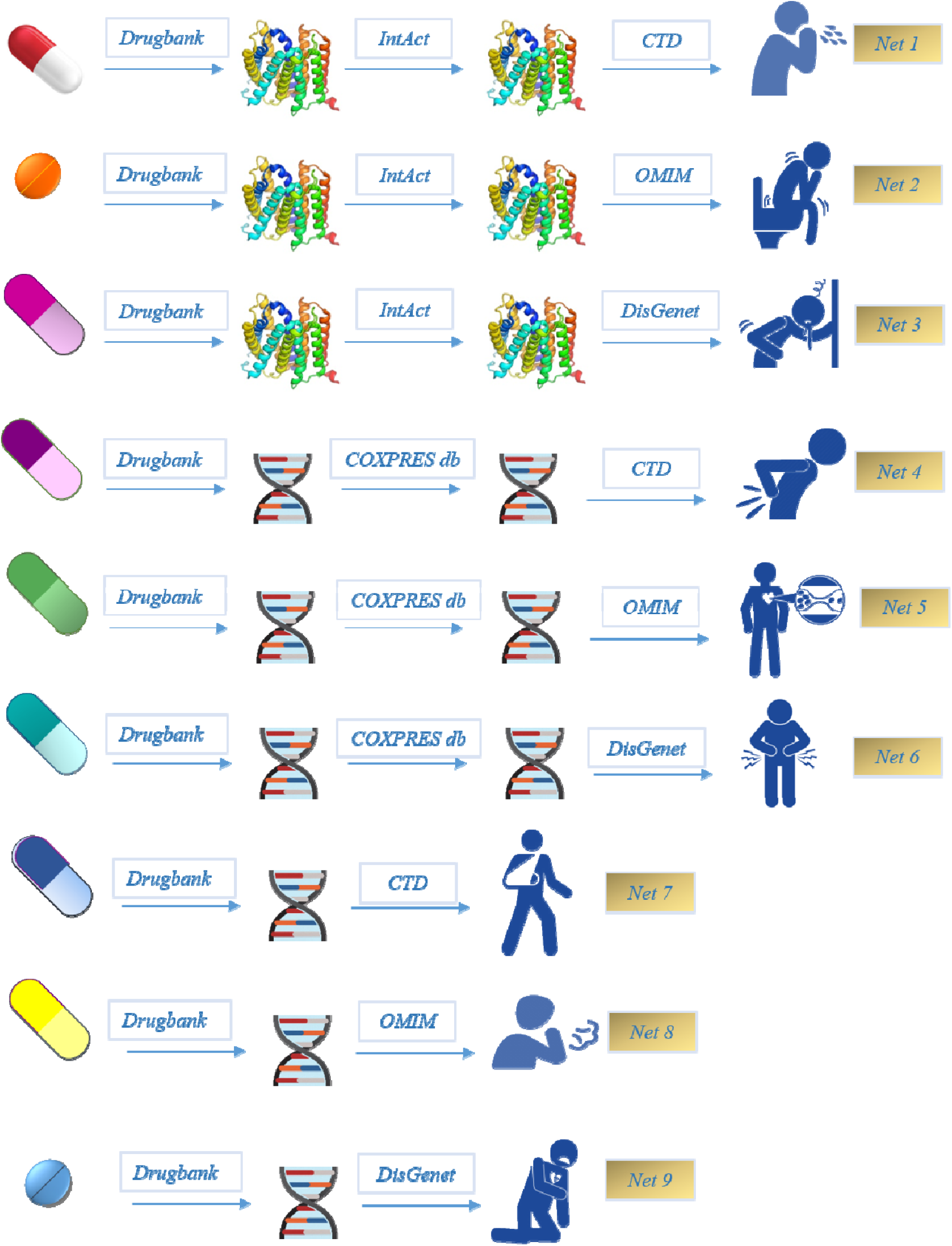
Schematic overwie(overview) of reconstructing nine new drug-disease networks.

### 2.3 Drug-disease association prediction

#### Encoding drug-disease networks as feature vectors

For each drug-disease pair, weights of its corresponding interaction in the reconstructed drug-disease networks were considered as features. Therefore, each drug-disease pair was encoded as a 9002Ddimentional feature vector.

#### Machine learning methods

We have used five different classifiers including naïve Bayes (NB), random forest (RF), logistic regression (LR), decision tree (DT) and support vector machine (SVM). The implementations of these classifiers in Weka [21] software package was used for drug-disease association prediction. Weka is a java based machine learning workbench that was developed for machine learning tasks. Also, we used 10-fold cross validation for evaluating the predicted drug-disease associations.

For evaluating the performance of RepCOOL, we used 4 different measures (Table 3). These measures are based on the following four basic terms:

**Table 3.** Measures for assessing prediction performance

|  |  |
| --- | --- |
| $Recall = \frac{TP}{TP + FN}$ | positive correctly predicted |
| $Precision = \frac{TP}{TP + FP}$ | Positive Predictive Value |
| $Accuracy = \frac{TP + TN}{TP + TN + FP + FN}$ | correctly predicted |
| $F - measure = \frac{2 \times Precision \times Sensitivity}{Precision + Sensitivity}$ | The harmonic mean of sensitivity and specificity |

True positive (TP): the number of drug-disease associations, which have been correctly predicted.

True negative (TN): the number of drug-disease pairs, which have been correctly predicted as non-associated.

False positive (FP): the number of unrelated drug-disease pairs, which have been incorrectly predicted as associations.

False negative (FN): the number of drug-disease associations, which have been incorrectly predicted as non-associations.

We also, used area under ROC curve (AUC) as another measure for assessing the proposed method.

#### Benchmark dataset

We used PREDICT [22], which is a well-known benchmark dataset in drug repositioning, to assess the strength of the proposed drug repositioning method. PREDICT dataset includes 1,834 interactions between 526 FDA approved drugs and 314 diseases.

## Results and discussion

### Performance evaluation of the proposed method

Figure 3. shows the performance of five classifiers on the PREDICT dataset in a 10-fold cross validation experiment. As it is evident, decision tree is the most sensitive classifier in detecting true drug-disease associations but random forest has the best performance in term of ROC. For all the classifiers recall (sensitivity) is in a satisfactory range, which shows the ability to detect true drug-disease associations. However, precision is relatively low for almost all classifiers, which can be a result of some true drug-disease associations that has not been discovered or reported yet.

**Figure 3:**
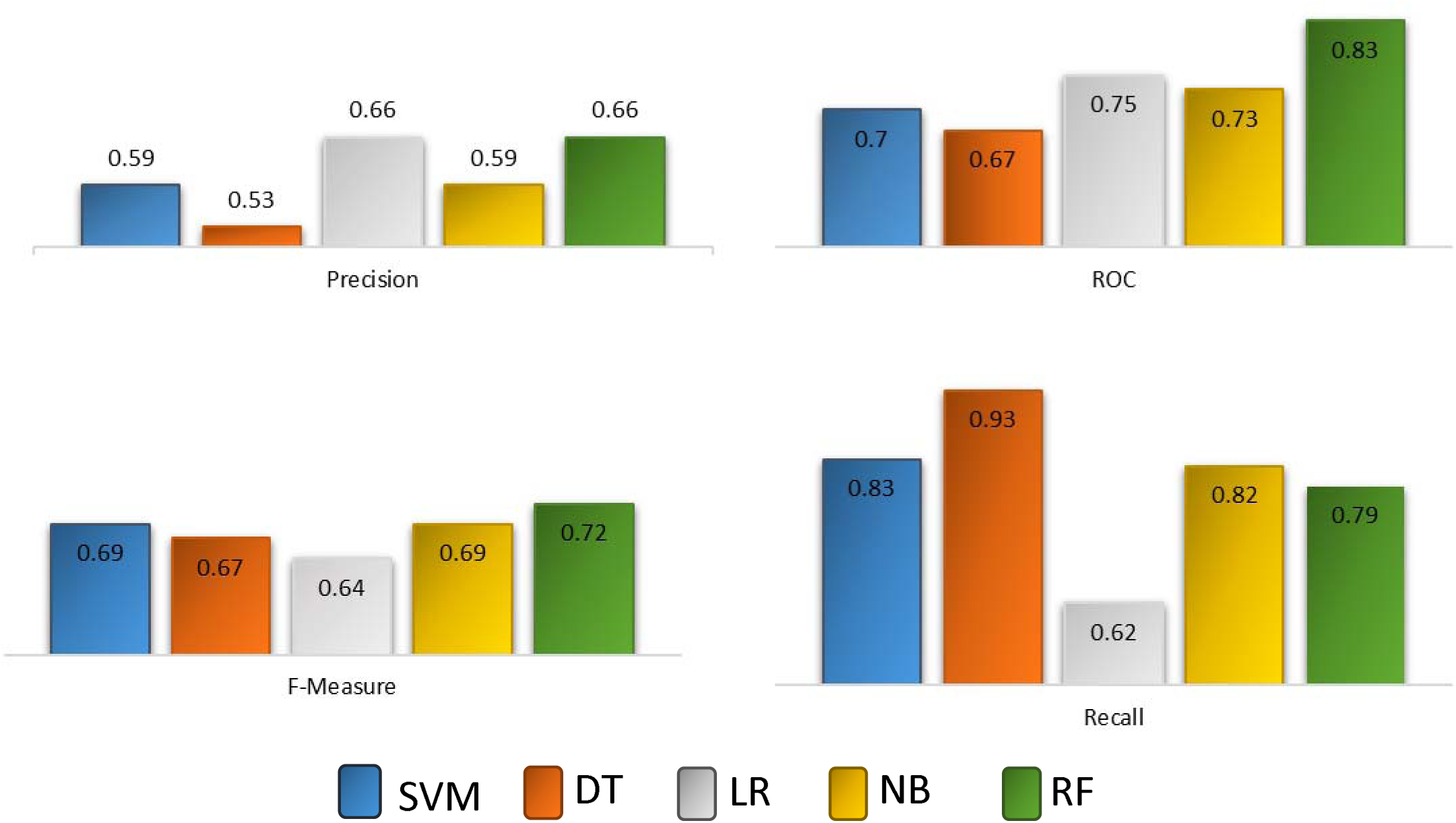
Performance of different classifiers in a 10-fold cross validation procedure in PRIDICT dataset. Classifiers include support vector machine (SVM), decision tree (DT), linear regression (LR), naïve Bayes (NB) and random forest (RF).

### Comparison with the other methods

Nearly all of the previously published studies only reported their AUC. As it is shown in figure 4 the highest AUC of the five classifiers is 0.83, which shows better performance than HGBI[23], LDB[24], TL-HGB[25] and Drug Net[24] methods on PREDICT dataset.

**Fig 4.**
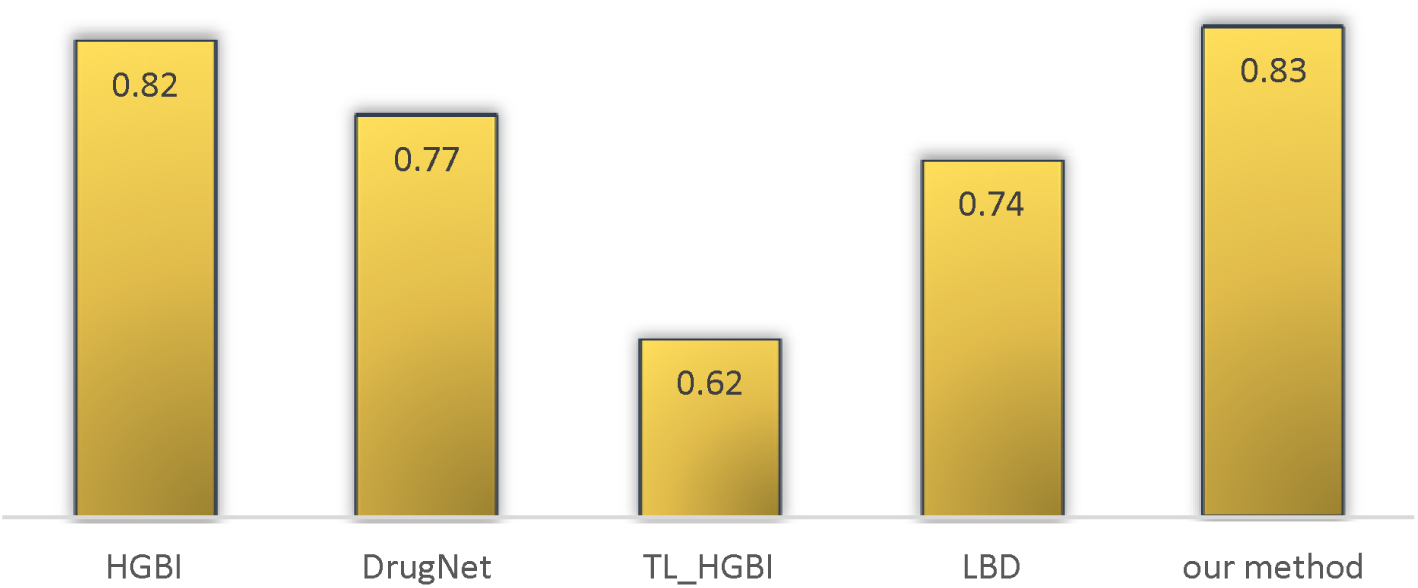
Performance comparison of RepCOOL with other methods in terms of AUC based on the obtained results in PREDICT dataset.

### New repurposed drugs for breast cancer

Information contained in RepoDB [26] was exploited to obtain a list of new repurposed drug for breast cancer. RepoDB includes a gold standard set of drug repositioning which have been failed or succeeded. The RepoDB dataset contains 6,677 approved, 2,754 terminated, 483 suspended and 648 withdrawn drug-disease interactions. Withdrawn and suspended drug-disease associations have annotation phase between phase 0 and phase 3. Therefore, these two types of drug-disease pairs have more potential to suggest a valid new drug repositioning rather than a random pair. Considering this fact, we have trained the five classifiers using the approved and terminated data. Figure 5 shows the training performance of the classifiers. Then, the best performing classifier, according to the approved and terminated data, was used to predict new drugs for breast cancer. The most sensitive classifier, which was random forest (it detected 2,283 true drug-disease interactions out of 2,292), was used to do this end.

**Figure 5:**
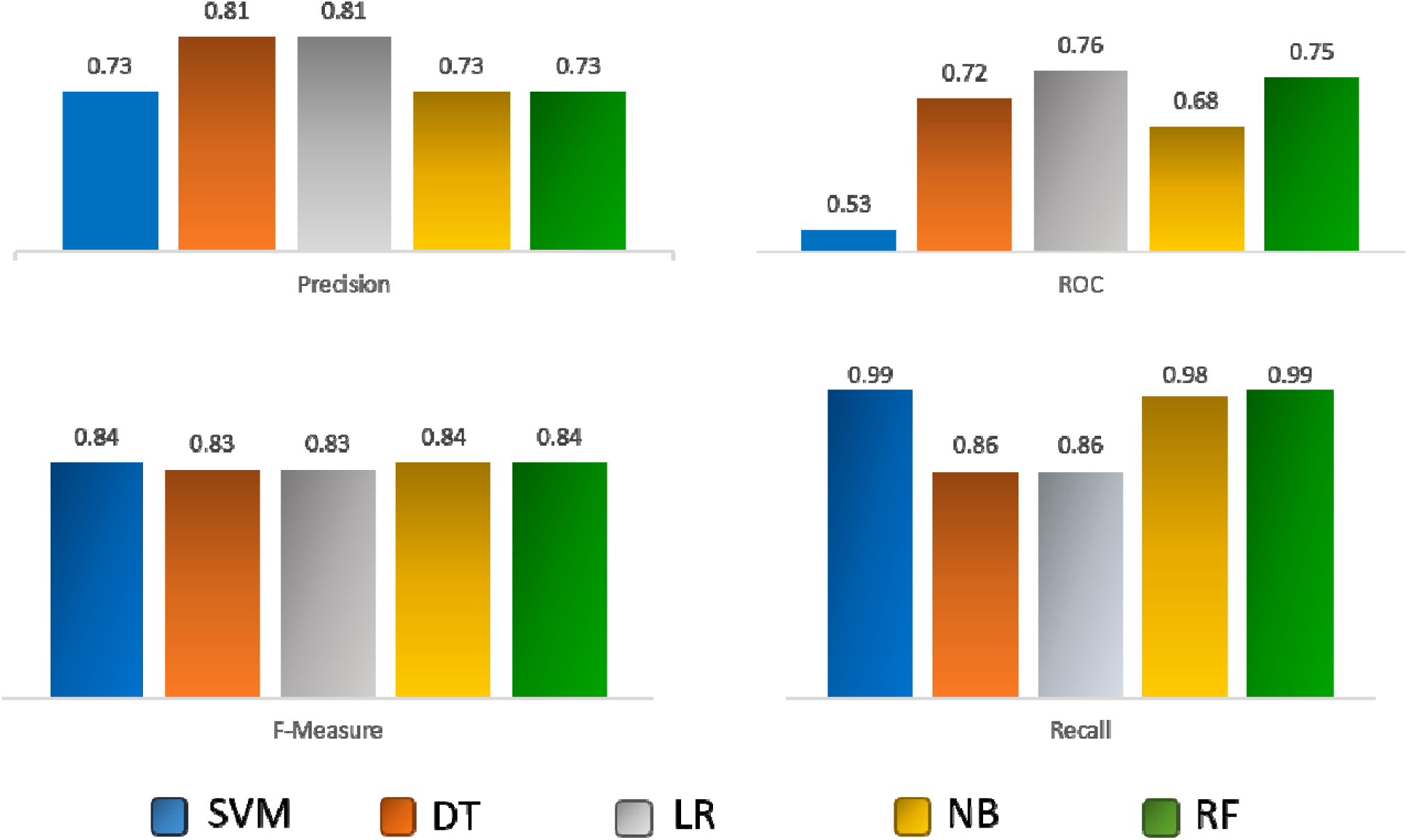
Performance of different classifiers in a 10-fold cross validation procedure in repODB dataset. Classifiers include support vector machine (SVM), decision tree (DT), linear regression (LR), naïve Bayes (NB) and random forest (RF).

Using this classifier, four new drugs have been repurposed for breast cancer stage II. Table.3 shows the chemical structures for these drugs and a brief description for each one.

### Analyzing the structural similarity between the four new repurposed drugs and previously FDA-approved drugs for breast cancer

We also did a structural similarity analysis between the repurposed drugs and 10 FDA-approved drugs for breast cancer including 5-FU, Abemaciclib, Verzeino, Taxotere (docetaxol), danazol, Pamidronate Disodium, Trastuzumab, Tamoxifen, Doxorubicin, Paclitaxel, Capecitabine, Dutasteride, Olaparib, Afinitor. Figure 6 shows the results of the structural similarity analysis. Structural similarity was computed based on 3,014 structural features that was extracted using Dragon tool [27]. Figure 5.a compares the structures of the drugs via a distance matrix, and figure 5.b represents the correlation matrix of the structures that was computed using Pearson correlation coefficient (PCC). Also, figure 5.c depicts the dendrogram of 14 drugs based on the obtained distance matrix. According to this dendrogram, we can see four distinct clusters: cluter1= {Danazol}, cluster2 = {Doxorubicin, Dutasteride, Taxotere, Abemaciclib, Paclitaxel, Olaparib, Trastuzumab, 5FU, Verzeino}, cluster3={Afinitor} and cluster4 = {Pamidronate Disodium, Capecitabine, Tamoxifen}. As it evident Paclitaxel, Doxorubicin and Tamoxifen have the most structural similarity with Abemacilib (PCC= 100), Dutasteride (PCC=100) and Capecitabine (PCC=94), respectively. Also 5FU and Verzeino are the two most similar FDA-approved drugs to the Trastuzumab with PCC of 99.

**Figure 6.**
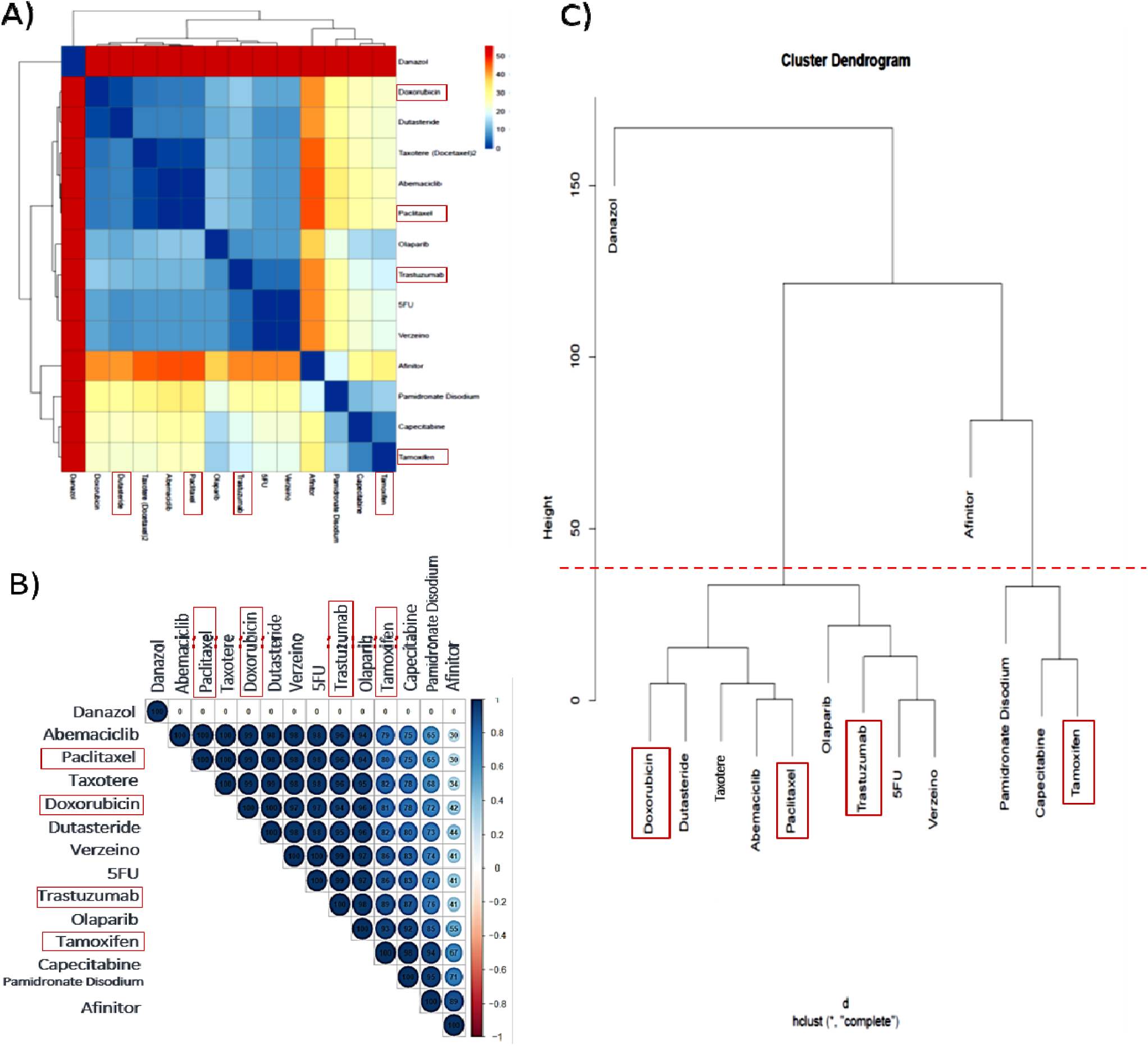
Structural relationship between the repurposed (highlighted by rectangles) and *FDA-approved drugs* for the treatment of breast cancer. (**A)** Heat map of the merged repurposed and *FDA-approved drugs* based on distance matrix. (**B)** Cluster dendrogram of repurposed and *FDA-approved drugs* based on distance matrix. **(C)** Heat map of repurposed and *FDA-approved drugs* based on correlation matrix. The highest and the lowest structural correlation are indicated in blue and red, respectively.

**Table 3.**
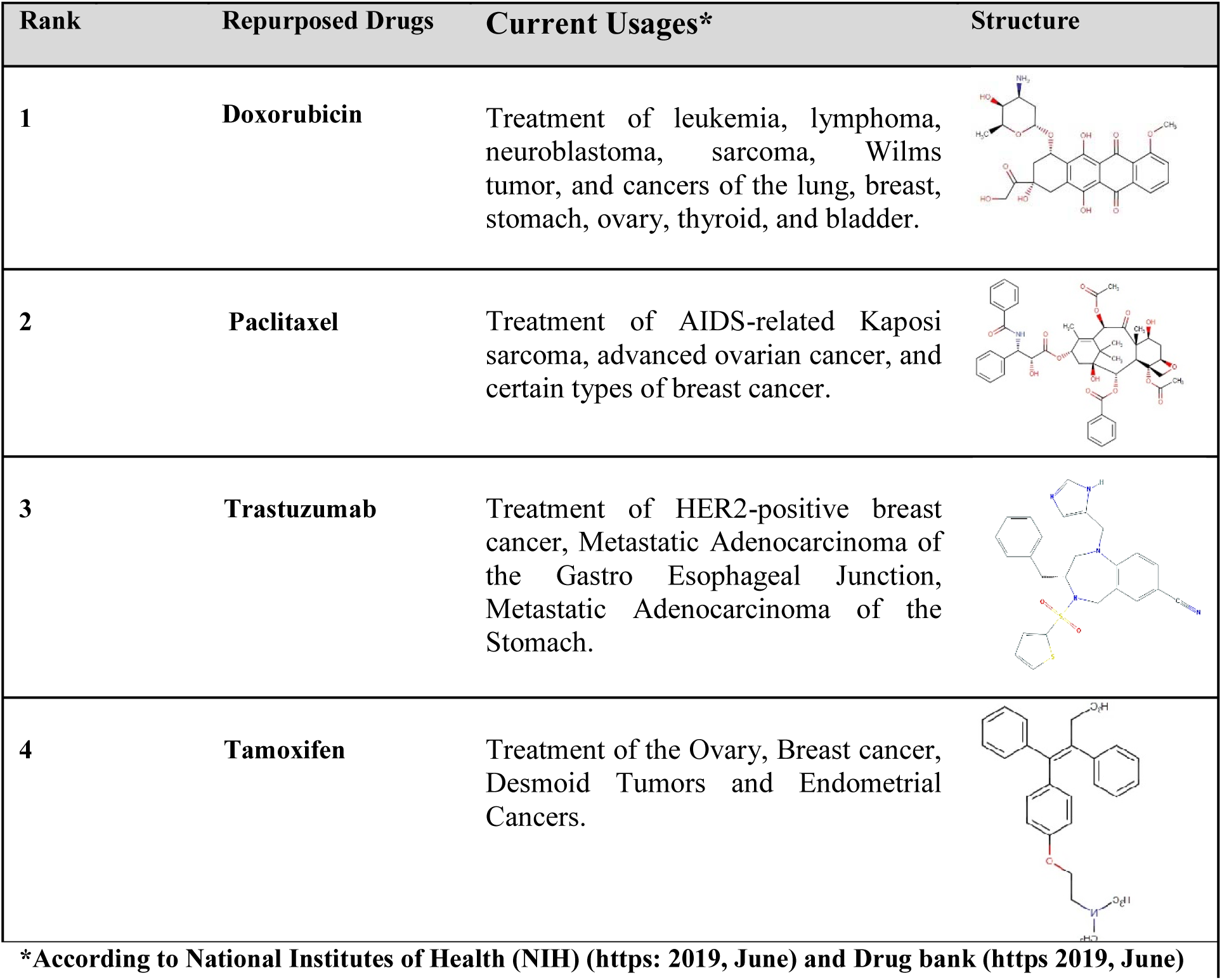
Summary of function and structure of the repurposed drugs for breast cancer.

| Rank | Repurposed Drugs | Current Usages* | Structure |
| --- | --- | --- | --- |
| 1 | **Doxorubicin** | Treatment of leukemia, lymphoma, neuroblastoma, sarcoma, Wilms tumor, and cancers of the lung, breast, stomach, ovary, thyroid, and bladder. |  |
| 2 | **Paclitaxel** | Treatment of AIDS-related Kaposi sarcoma, advanced ovarian cancer, and certain types of breast cancer. |  |
| 3 | **Trastuzumab** | Treatment of HER2-positive breast cancer, Metastatic Adenocarcinoma of the Gastro Esophageal Junction, Metastatic Adenocarcinoma of the Stomach. |  |
| 4 | **Tamoxifen** | Treatment of the Ovary, Breast cancer, Desmoid Tumors and Endometrial Cancers. |  |

\*According to National Institutes of Health (NIH) ([https: 2019, June](https://www.nih.gov/)) and Drug bank ([https 2019, June](https://www.drugbank.ca/))

### MTT Assay

An MTT assay was also done to assess the effectiveness of the repurposed drugs (figure 7). According to our limitations we did the MTT assay only for tamoxifen. Human cell line BT474 was cultured in recommended media in the presence of 10% fetal bovine serum (FBS) and penicillin-streptomycin antibiotics. Cell viability was characterized using a standard colorimetric MTT (3-4,5-dimethylthiazol-2-yl-2, 5-diphenyl-tetrazolium bromide) reduction assay. Briefly, 6000 cells were plated in each well of the 96-well plates with 100 µL medium which includes 10% serum. After 24-hour incubation, the cell was treated with several concentration of tamoxifen (0-100µM). After 48-hour the MTT reagent (5 mg/ml in PBS) was added to each well, followed by incubation for four hours at 37 c with 5% CO2. After the incubation, the MTT crystals in each well were solubilized in 100µl DMSO incubation for 20 min at 25 c, and the absorbance was read at 490 nm with an ELISA reader.

**Figure 7.**
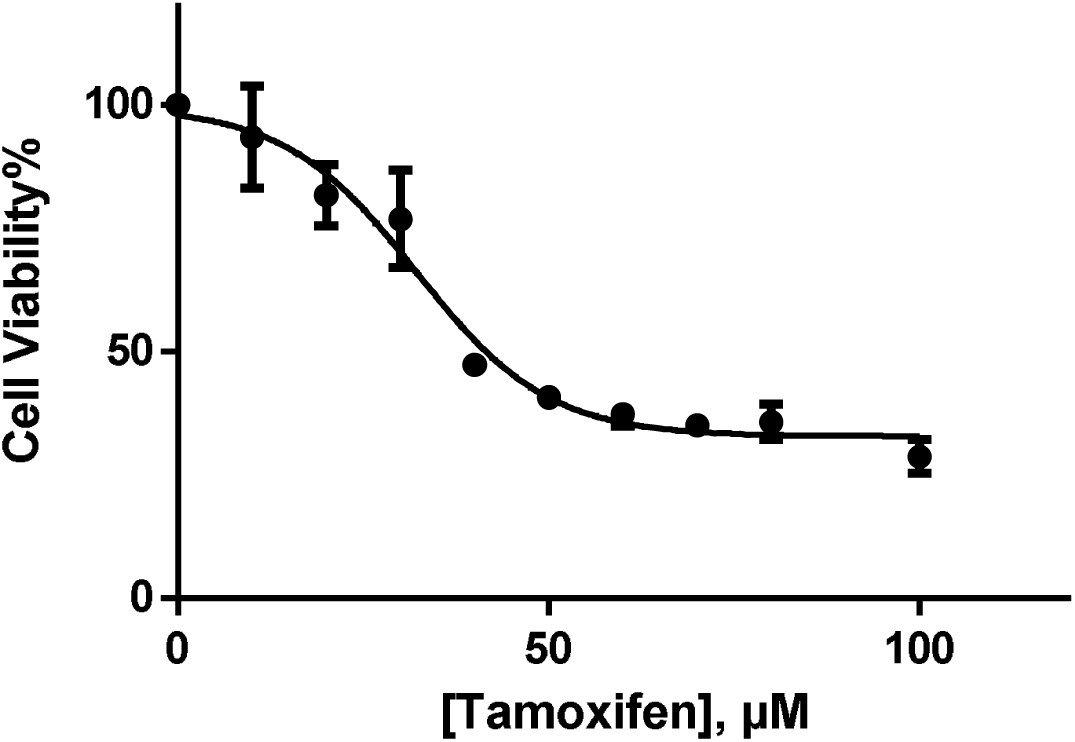
IC_50_ plot. Inhibitory effect of different concentrations of tamoxifen on growth of BT474 cells. The vertical axis determines the absorbance and horizontal axis shows the tamoxifen concentration.

Cell survival following treatment with tamoxifen was measured using an MTT assay to evaluate the effect of the growth inhibition on the breast cancer stage II, HER2 cell line. The figure 6 shows the absorption of tamoxifen in live cells at 490 nm. According to the obtained result, the half maximal inhibitory concentration (IC50) of tamoxifen was 32.13 µM.

### Conclusion

In this study a network based approach was exploited for drug repositioning using heterogeneous biological and chemical information. Results show the strength of the proposed method in detecting true drug-disease relationships. RepCOOL suggested four new drugs for breast cancer stage II namely Doxorubicin, Paclitaxel, Trastuzumab and Tamoxifen. Structural analysis showed high structural similarity of this four drugs to the current FDA-approved drugs for breast cancer stage II. In addition, we did an MTT assay for one of the suggested drugs (Tomoxifen), which had IC50 of 32.13 µM.

## Abbreviations

FDA: Food and Drug Administration
DRGN: drug-gene interaction network
DIGN: disease-gene interaction network
PPIN: protein-protein interaction network
GCN: gene co-expression network
CTD: Comparative Toxic genomics Database
OMIM: Online Mendelian Inheritance in Man
GDAs: gene–disease associations
NB: naïve Bayes
RF: random forest
LR: logistic regression
DT: decision tree
SVM: support vector machine
TP: True positive
TN: True negative
FP: False positive
FN: False negative
ROC: Receiver Operator Characteristics
AUC: Area Under the Curve
PCC: Pearson correlation coefficient
MTT: 3-(4,5-dimethylthiazol-2-yl)-2,5-diphenyltetrazolium bromide
FBS: fetal bovine serum
DMSO: dimethyl sulfoxide
IC50: concentration giving half-maximal inhibition

## Acknowledgements

No applicable.

## Declarations

## Availability of data and material

No applicable.

## Funding

No applicable.

## Authors ‘ contributions

LL, ML and TD designed the algorithm for data curation. YZ, LL, XZ, CM and ZW implemented the front end and back end of the tool. ZW, LL and JL wrote the manuscript and discussed the results. All authors read and approved the final manuscript.

## Ethics approval and consent to participate

Not applicable.

## Consent for publication

Not applicable.

## Competing interests

The authors declare that they have no competing interests.

